# Statistical Model of Motor Evoked Potentials for Simulation of Transcranial Magnetic and Electric Stimulation

**DOI:** 10.1101/406777

**Authors:** Stefan M. Goetz, S. M. Madhi Alavi, Zhi-De Deng, Angel V. Peterchev

## Abstract

Motor evoked potentials (MEPs) are widely used for biomarkers and dose individualization in transcranial stimulation. The large variability of MEPs requires sophisticated methods of analysis to extract information fast and correctly. However, models of MEPs that represent their characteristic features are lacking. This work presents a statistical model that can simulate long sequences of individualized MEP amplitude data with properties matching experimental observations. The MEP model includes three sources of trial-to-trial variability to mimic excitability fluctuations, variability in the neural and muscular pathways, and physiological and measurement noise. It also generates virtual human subject data from statistics of population variability. All parameters are extracted as statistical distributions from experimental data from the literature. The model exhibits previously described features, such as stimulusintensity-dependent MEP amplitude distributions, including bimodal ones. The model can generate long sequences of test data for individual subjects with specified parameters or for subjects from a virtual population. The presented MEP model is the most detailed to date and can be used for the development and implementation of dosing and biomarker estimation algorithms for transcranial stimulation.

## I. Introduction

Direct neural responses are important for detecting the effects of transcranial magnetic stimulation (TMS) and suprathreshold transcranial electrical stimulation (TES). Two measures presently used to characterize cortical effects and to calibrate stimulation intensity are the phosphene and motor thresholds. Direct suprathreshold stimulation of the visual cortex produces phosphenes, which the subject experiences as artifacts in his or her vision, but are difficult to reliably quantify [1–5]. In contrast, stimulation in the motor cortex with sufficient strength can initiate motor evoked potentials (MEP) that can be recorded as a short wave of electric activity through electromyography (EMG) at the corresponding peripheral muscle. MEPs show a wide dynamic range from microvolts to millivolts, allowing for good quantification [6–9]. The immediate responses of other brain targets are presently only visible in electroencephalography at the cost of higher acquisition effort, more pronounced noise, and yet poorer interpretability [10–12].

Due to easily accessible and quantifiable MEPs, the primary motor cortex is one of the most important targets for brain stimulation. Noninvasive stimulation of the primary motor cortex is used for the diagnosis and localization of motor lesions [13]. Furthermore, the primary motor cortex is a preferred model for studying the neurophysiology, biophysics of brain stimulation, and development of novel technology [14–16]. According to safety guidelines of repetitive TMS, the motor threshold is the reference of individual dosage also for other brain targets that are silent [17]. Individualization of stimulation relative to the motor threshold contributes to ensuring safe ranges of the stimulation parameters and maximizing effect size, repeatability, as well as translatability of the results [2, 18–20]. Importantly, the FDA requires the determination of the motor threshold before any therapeutic intervention with repetitive TMS, such as the treatment of major depression [21, 22].

MEPs in response to any brain stimulation technique, however, show complicated dependencies on the stimulus parameters and are highly variable [23–32]. Identical consecutive stimuli can evoke different responses, from undetectable signal to one with saturated amplitude. This variability is not the result of only additive measurement noise, e.g., from amplification, but includes neural variability processes from various sources, such as rapid excitability fluctuations of the targeted neurons due to incoming endogenous signals [33–39]. These characteristics can vary both across subjects as well as within subjects, e.g., due to endogenous brain state or exogenous interventions [40–42]. Thus, the variability of MEPs may in itself be an interesting biomarker of the underlying neural circuits, and therefore should be correctly incorporated in MEP models.

The complexity of MEP responses further requires appropriate mathematical and statistical methods to estimate accurately and efficiently parameters such as target hot spot, motor threshold, or neural recruitment input–output (IO) curves [43–50]. In order to minimize the number of required stimulation pulses, estimation methods ideally analyze MEP responses online and adaptively adjust parameters based on the previous observation, e.g., at which stimulation strength to stimulate next for maximum information extraction [51]. The development and evaluation of such methods require intensive testing under realistic conditions. Such testing requires many subjects, test conditions, and repeated trials, typically well above a thousand [46, 47, 52, 53]. This is typically not practical in an experimental study, especially for early stages of development and for comparison of the fundamental properties of various estimation algorithms. Previously, due to the lack of independent realistic models, testing and validation of brain stimulation methods against models was sometimes done with the very model that the method estimates internally. However, such testing against internal models becomes cyclic and self-fulfilling such that the test only confirms its own model assumptions. If a method is well adjusted to specific data or test models but is challenged in real experiments, it typically suffers from so-called model bias. Thus, experiments in subjects are vital through the course of development and reveal bias of the methods towards the simulation model. However, they cannot satisfy the vast testing needs at the beginning of and during design [54]. Furthermore, progress is hampered as many theoreticians from mathematical, physical, or engineering disciplines do not have access to experimental setups or raw data.

In this paper, we present a realistic model of MEP amplitude for brain stimulation, especially TMS and TES, that incorporates the intra-individual variability as well as inter-individual spread of features of MEPs. The model can generate virtual subjects, which reflect the spread of MEP features within the population. For a given subject, the model can generate MEPs in dependence of the stimulation strength that show trial-to-trial variability with experimentally established statistical distribution. A software implementation of the model is available online for the research community to use.

## II. Methods

We aim to construct a generative statistical model of MEP amplitude as a function of stimulation strength in the primary motor cortex. The model requirement is to incorporate both intra-individual variability, i.e., trial-to-trial response fluctuation, and inter-subject variability, i.e., the individual neural recruitment features.

The proposed model builds upon a statistically validated structure we published previously [42]. For a normalized input stimulation intensity, *x*, and output peak-to-peak MEP voltage, *y*, the model consists of five parameters to describe the MEP distribution. We tested each model parameter for its inter-individual distribution with initial distribution screening based on Akaike information and subsequent Shapiro–Wilk test in case of normal and lognormal distributions. Subsequently, the parameters underwent log-likelihood estimation to derive more general statistical distributions. Based on the basic model and individual parameters, we generalized the parameters to population distributions that allow the generation of new virtual subjects based on the statistical distributions derived from a previously analyzed subject population of 12 subjects for 60 µs monophasic cTMS pulses [42].

In addition, the model incorporates three variability parameters to describe inter-individual variability, intra-individual trial-to-trial variability, as well as physiological and measurement noise. The inter-individual multiplicative output-side or *y* variability and additive input-side *x* variability are taken from the literature [42]. We extended this model with additive *y* variability to represent measurement noise, such as electrode, environmental, and amplifier noise, to the physiological model and derived an accurate distribution from recordings. Importantly, the additive *y* variability is, in contrast to common expectations, non-Gaussian for MEPs. Even though it may have Gaussian origin in the continuous EMG recording, the peak-to-peak MEP amplitude is highly sensitive to only the largest spike within the short range of an apex of an MEP wave [55]. Therefore, it amplifies the impact of outliers and converts well-studied canonical noise distributions such as Gaussian noise into extreme value distributions with heavily skewed tail. The additive noise affects only the low-amplitude sections of the IO curve and leads to the low-side plateau.

To estimate the distribution of the additive *y* variability, we used sections of the original EMG recordings. These EMG recordings were acquired with a commercial amplifier (BIOAMP-4, SA Instrumentation Co., San Diego, CA; bandwidth 30 Hz to 1 kHz) and neonatal ECG electrodes with 25 mm total and approximately 10 mm active diameter (Kendall Kittycat, Medtronic Co., Minneapolis, MN). We extracted more than 2900 recording epochs between stimuli at least 6–8 s after a previous TMS pulse and before the next one. The epochs had the same duration as the windows used for MEP peak-to-peak amplitude detection (17.5 ms duration) and were subjected to identical filtering (fourth-order Butterworth low-pass filter with 1 kHz cut off). Whereas the raw sample-to-sample noise may be Gaussian (*p*(*W >* 0.99) *>* 0.08), the noise undergoes the same peak-to-peak amplitude extraction as MEPs, which transforms it to a general extreme-value distribution.

### A. Model Specification

Fig. 1A shows a diagram of the MEP model structure. The peak-to-peak MEP amplitude, *V*_pp_, in volts as a function of stimulation pulse strength, *x*, in normalized units, typically portion of maximum stimulator output in the range of 0 to 1 (or, equivalently, 0 to 100%), is described by

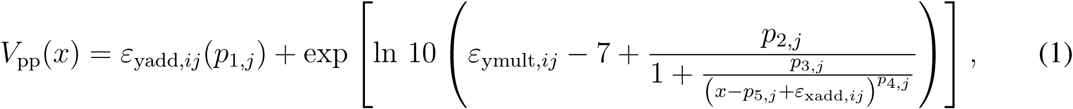

**Fig. 1.**
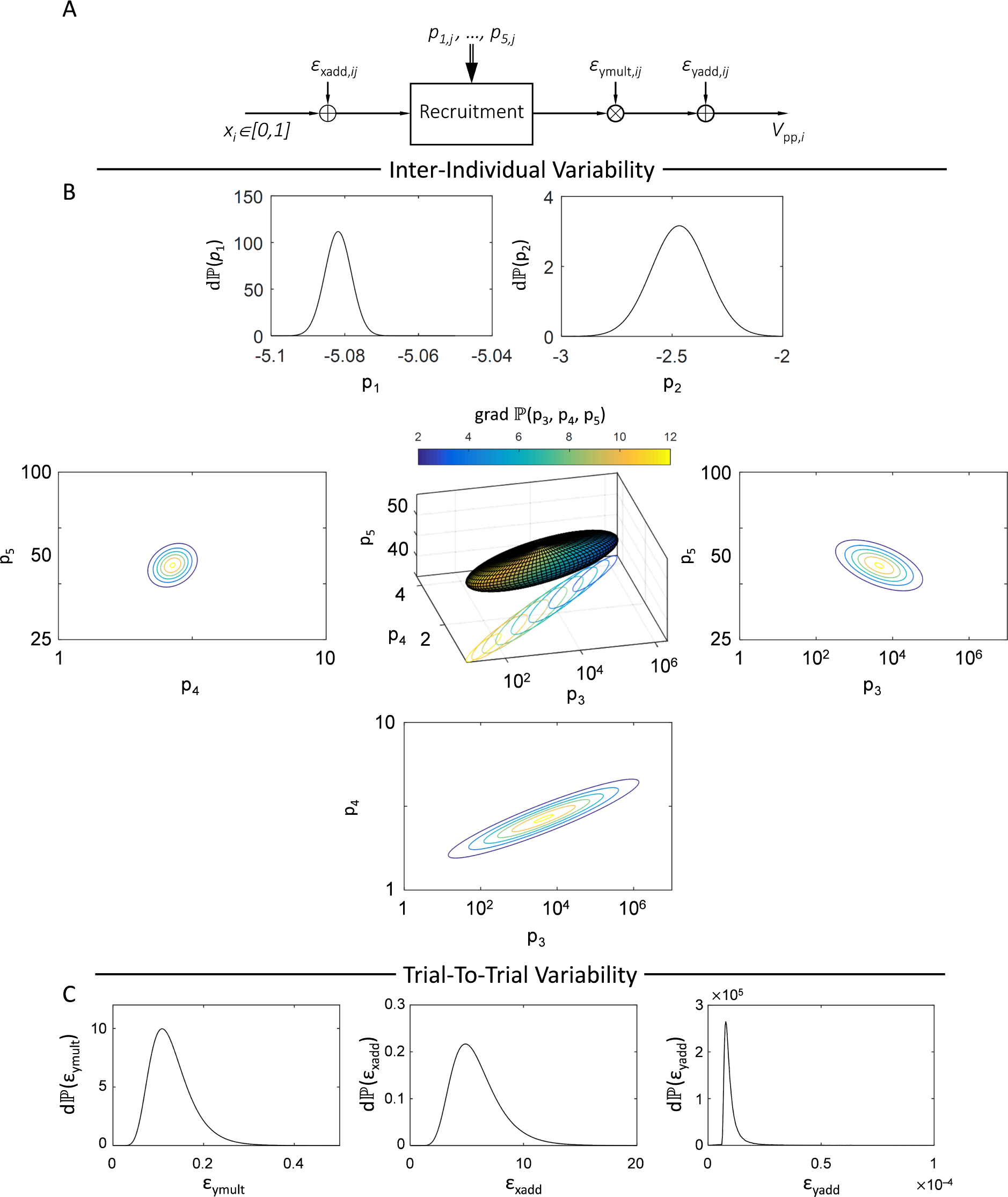
(A) MEP amplitude model structure. Distribution of (B) inter-individual (*p*_1,*i*_ to *p*_5,*i*_) and (C) intra-individual (*ε*_ymult,*ij*_, *ε*_xadd,*ij*_, and *ε*_yadd,*ij*_) parameters to represent, respectively, the spread of MEP features in the subject population and trial-to-trial variability.

where *p*_1,*j*_ to *p*_5,*j*_ are the individual recruitment parameters for subject *j*.

For the population of our experimental study [42], *p*_1,*j*_ (*p*(*W >* 0.87) *>* 0.161) and *p*_2,*j*_ (*p*(*W >* 0.97) *>* 0.90) show normal distributions, whereas *p*_3,*j*_, *p*_4,*j*_, and *p*_5,*j*_ are exclusively positive and exhibit approximately lognormal distributions (*p*(*W <* 0.43) *<* 0.001, *p*(*W >* 0.92) *>* 0.45, and *p*(*W >* 0.94) *>* 0.63, respectively). Parameters *p*_3,*j*_, *p*_4,*j*_, and *p*_5,*j*_ are correlated (*p*(*r*_34_ *>* 0.97) *<* 0.001, *p*(*r*_35_ *< -*0.87) *<* 0.005, and *p*(*r*_45_ *< -*0.82) *>* 0.012), whereas other parameter pairs are not. The individual recruitment parameters show the following normal and lognormal distributions

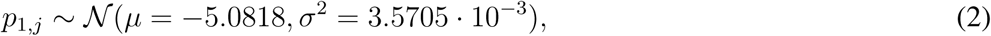

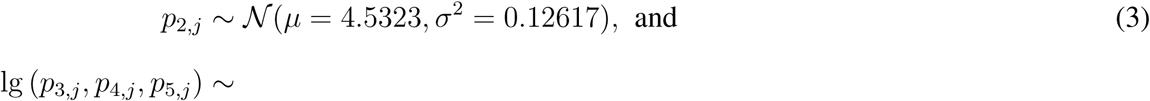

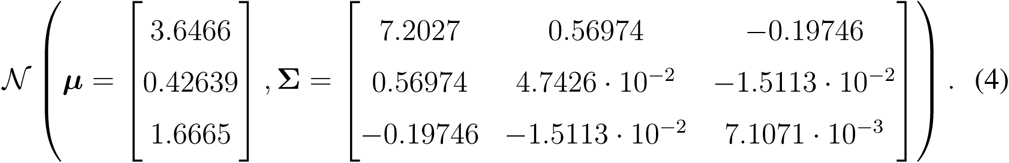

Fig. 1B illustrates the inter-individual distributions of *p*_1,*j*_,…, *p*_5,*j*_. Parameters *p*_1,*j*_ and *p*_2,*j*_ are shown as one-dimensional density function plots. Since parameters *p*_3,*j*_, *p*_4,*j*_, and *p*_5,*j*_ are highly correlated, they are depicted as three-dimensional isoquant surface and as cross sections in all three dimensions through their expectancy value, 𝔼(*p*_3_, *p*_4_, *p*_5_).

The intra-individual variability sources *ε*_ymult,*ij*_ and *ε*_xadd,*ij*_ as well as the additive noise *ε*_yadd,*ij*_, which may be dominated by measurement noise, vary between pulses *i* and subjects *j* along distributions with subject-dependent parameters. The former two show lognormal distributions (*p*(*W >* 0.98) *>* 0.97 and *p*(*W >* 0.90) *>* 0.31), whereas the latter follows an extreme-value distribution as discussed above. The three variability terms can accordingly be sampled from

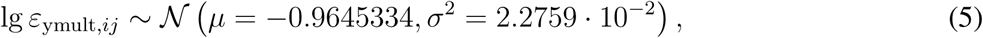

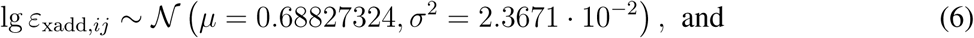

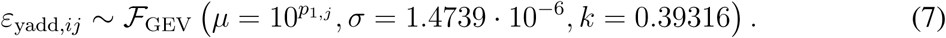

The common logarithm is denoted by lg(*x*) *≡* log_10_ *x*. The generalized extreme value distri bution *ℱ* _GEV_ follows [56, 57]

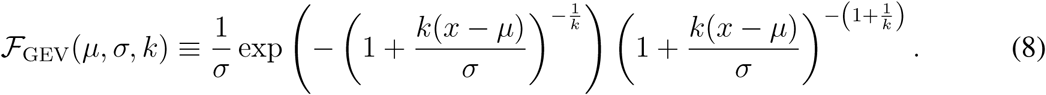

Fig. 1C displays the distributions of the trial-to-trial variability fluctuations *ε*_ymult,*ij*_, *ε*_xadd,*ij*_, and *ε*_yadd,*ij*_.

## III. Results

To generate a virtual subject with this model, one has to assign parameters *p*_1_ through *p*_5_. These parameters can be assigned random values according to the distributions in Eqs. (2)– (4) to represent a random member of the population. Alternatively, one can determine the parameter values corresponding to a specific real subject of interest, e.g., by fitting the model to experimentally measured responses. With the individual parameters set, MEP amplitudes in response to stimuli with certain stimulation strength require the random generation of trial-to-trial variability and noise parameters *ε*_ymult,*ij*_, *ε*_xadd,*ij*_, and *ε*_yadd,*ij*_ before each stimulus according to the distributions given by Eqs. (5)–(7).

Although the model describes MEP features only by few parameters and their statistical distribution, the model presents high-level characteristics reported in the literature. For example, Fig. 2A plots a model-generated IO curve, displaying previously known features, such as heteroscedastic spread of samples with a relatively narrow right-skewed spread at low stimulation strengths, a large spread around the motor threshold and the slope section of the IO curve, and a moderate symmetric to left-skewed spread at saturation [24, 50]. Furthermore, under certain conditions, the model presents bimodal distribution of MEPs at fixed stimulation strength in some subjects close to the threshold as shown in Fig. 2B [33].

**Fig. 2.**
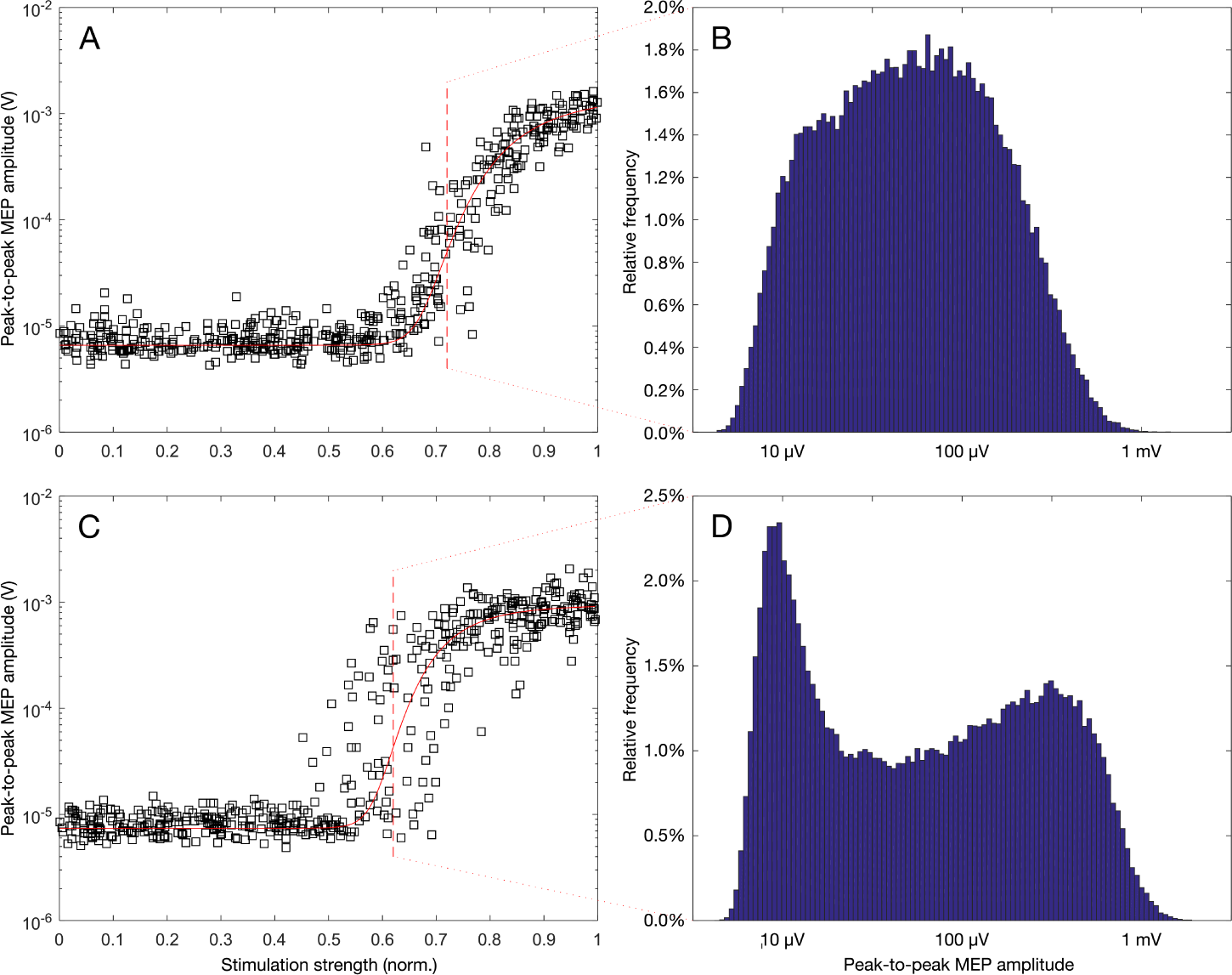
Model-generated IO curves of two virtual subjects, including 500 uniformly distributed samples and corresponding average sigmoid curve (A and C), and 100,000 samples at a single stimulation strength point within the curve slope (B and D). These examples illustrate previously described MEP features such as approximately lognormal distribution around the threshold (B) and locally bimodal distribution around the threshold (D).

## IV. Conclusions

We presented a statistical model and its parameters that generate MEP amplitude data as a function of stimulation strength with realistic variability distribution. The distributions of all model parameters are estimated for a published TMS study, allowing the generation of virtual subjects that represent typical population statistics. The model incorporates intra-and inter-individual variability and manifests previously described statistical features of MEPs. Although it is a phenomenological model, it allows the analysis and extraction of MEP metrics, such as motor-threshold values, as well as the study of the influence of certain parameters on the IO curve. Furthermore, the model enables design and realistic testing of methods for brain stimulation, such as threshold and IO curve estimation, including consideration and assessment of their variability. The stimulation strength can be scaled to practically any available supra-threshold brain stimulation device, though differences in parameters, such as the intra-individual parameters, might arise and should be studied in the future. Finally, this model assumes that all of the MEPs are independent; future work should explore the pulse-to-pulse relationship as a function of inter-pulse interval and pulse-train duration.

## Code Availability

An implementation of the model in MATLAB accompanied by instructions for its use is available at https://github.com/sgoetzduke/Statistical-MEP-Model. This model can be freely used and enhanced by the research community.

## Acknowledgments

This work was funded by the NIH under grants R01NS088674 and RF1MH114268; Brain & Behavior Research Foundation NARSAD Awards 22796 and 26161; and the North Carolina Biotechnology Center Grant 2016-CFG-8004. Z.-D. Deng is support by the National Institute of Mental Health Intramural Research Program.

The authors are inventors of brain stimulation technology not related to the matter of this paper. There are no relevant conflicts of interest.

